# Stepwise Proteinase K Digestion Improves Genomic DNA Yield and Purity from Goat Semen: A Comparative Pilot Study of Five Extraction Protocols

**DOI:** 10.64898/2026.10.11.758327

**Authors:** Sharadindu Shil, Ankur Singha, Soumita Paul, Ashish Khatua, Shankari Prasad Dutta, Tarpan Mondal, Jayanta Chowdhury, Ayan Mukherjee, Anusuya Debnath, Ajoy Mandal

**Affiliations:** Centre for Laboratory Animal Research and Training (CLART), West Bengal Livestock Development Corporation Limited (WBLDCL), Government of West Bengal, Kalyani, Nadia, West Bengal, India; Department of Life Sciences, Garden City University, Bangalore, India; Department of Biotechnology, Brainware University, Kolkata, West Bengal, India; ICAR-National Dairy Research Institute, Eastern Regional Station, A-12 Block, Kalyani, Nadia, West Bengal 741235, India

**Keywords:** artificial insemination, Black Bengal, chromatin, germplasm, polymerase chain reaction, spermatozoa

## Abstract

Genomic DNA of high quality is a prerequisite for molecular genetic analyses, but extraction from goat semen is difficult because sperm chromatin is highly compacted and seminal plasma contains inhibitors. This pilot study aimed to identify a reliable and cost-effective method for isolating genomic DNA from goat semen. Five extraction protocols were compared: the unmodified blood protocol (Protocol B), two modifications of a sperm protocol (Modified Protocols 1 and 2) and a stool protocol (Protocol BS) of a commercial spin-column kit (Kit A), and a second commercial silica membrane kit (Kit B). Three technical replicates per protocol were prepared from one frozen semen straw of a Black Bengal buck (one replicate for Protocol B). DNA concentration, purity (absorbance ratios at 260 and 280 nm, A_260_/A_280_, and at 260 and 230 nm, A_260_/A_230_), cost per sample and amplification of the mitochondrial hypervariable region 1 (HVR1) by polymerase chain reaction (PCR) were measured. Modified Protocol 2, which used 100 µL of semen and two 1.5-h proteinase K incubations at 55 °C, gave the highest normalised yield (mean ± standard deviation, 55.9 ± 30.4 ng µL^-1^ semen), the best purity (A_260_/A_280_, 1.78; A_260_/A_230_, 2.06) and the lowest cost per ng of DNA (INR 0.073 ± 0.037), and its extracts amplified successfully. The Kruskal–Wallis test for normalised yield was not significant (*p* = 0.075), so the differences in yield are exploratory. Because only one animal was sampled, the protocol must be confirmed with biological replicates from several bucks before wider use.

## INTRODUCTION

Goats (*Capra hircus* L.) are among the most important livestock species in tropical and subtropical regions and contribute to rural livelihoods, nutritional security and sustainable agricultural systems. Their adaptability to harsh environments, tolerance of disease and ability to thrive under low-input management make them indispensable in resource-constrained farming systems (FAO, 2020). Genetic improvement programmes for milk yield, meat quality, fibre production, reproductive efficiency and disease resistance rely on molecular genetic tools, which in turn depend on high-quality genomic DNA. The integrity, purity and yield of the DNA determine the success of downstream applications such as polymerase chain reaction (PCR), genotyping, marker-assisted selection and next-generation sequencing (Trost et al., 2019).

DNA is traditionally isolated from blood or tissue, but these approaches are invasive, require animal handling and may not be feasible under field conditions. Semen is a practical and minimally invasive alternative source of genomic DNA, particularly in artificial insemination programmes and germplasm conservation (Amills et al., 2017). Modified silica-column protocols have been used to obtain genotyping-grade DNA from cryopreserved sperm in fish (Fopp-Bayat & Ciereszko, 2012), which shows that archived semen can serve as a DNA source. Efficient DNA extraction from semen is nevertheless challenging. Spermatozoa have highly condensed chromatin stabilised by protamines and disulphide cross-links, which resist conventional lysis (Ward & Coffey, 1991; Hutchison et al., 2017; Kumaresan et al., 2020). Seminal plasma also contains inhibitory proteins and enzymes that interfere with downstream enzymatic reactions. The available techniques, including phenol–chloroform extraction and commercial column-based kits, involve trade-offs between yield, purity, cost, safety and processing time (Birren et al., 1997; Sambrook & Russell, 2001; Kumari et al., 2019). Comparative data for goat semen are scarce (Silva et al., 2014).

The aim of this pilot study was to compare five DNA extraction methods, including two modifications of a commercial spin-column genomic DNA kit, for yield, purity, cost and PCR amplifiability, and to identify a reliable, cost-effective method for isolating genomic DNA from goat semen. The study adds a comparison that includes the cost per unit of DNA. It used three technical replicates from a single semen straw, so the results are preliminary optimisation data and a basis for later studies with biological replicates from several animals.

## MATERIALS AND METHODS

### Study site and samples

The study was carried out at the CLART Genomics and Molecular Biology Laboratory (GMBL), Kalyani, Nadia, West Bengal, India, in the month of September 2025. Frozen semen straws of Black Bengal goat from routine artificial insemination archive stock were processed under aseptic conditions. For each protocol, three independent technical replicates were prepared from aliquots of a single frozen straw (one buck). The results therefore reflect reproducibility within protocols and not variation among animals. Protocol B was evaluated in a single extraction (n = 1) because of reagent constraints. Its data are reported descriptively and were excluded from the inferential statistics. No live animals were handled and no procedures were performed on animals, so approval by an Institutional Animal Ethics Committee was not applicable.

### DNA extraction protocols

Five protocols were evaluated with two commercial genomic DNA kits, and the DNA was stored at –20 °C until use. Protocol B (unmodified blood protocol, Kit A): DNA was extracted from 250 µL of semen with the QIAamp DNA Mini Kit (Qiagen, Germany; Cat. No. 51304) according to the standard blood protocol of the manufacturer (Qiagen, 2020). Modified Protocol 1 (Kit A): 250 µL of semen was processed with the same kit following the sperm protocol of the manufacturer (Qiagen, 2020) with slight modifications; the semen was washed with Buffer 1, 300 µL of Buffer 2 and 40 µL of proteinase K were added, and the sample was incubated at 55 °C for 2 h and then processed according to the protocol. Modified Protocol 2 (Kit A): 100 µL of semen was washed with Buffer 1, 300 µL of Buffer 2 and 10 µL of proteinase K were added, and the sample was incubated at 55 °C for 1.5 h; a second 10-µL aliquot of proteinase K was then added and the sample was incubated for a further 1.5 h at 55 °C. Buffer AL (200 µL) was then added and the sample was processed according to the protocol. Kit B: 200 µL of semen was processed with the PureLink Genomic DNA Mini Kit (Invitrogen, Germany; Cat. No. K182001) according to the recommendations of the manufacturer (Thermo Fisher Scientific, 2007). Protocol BS (Kit A-S, stool protocol): 200 µL of semen was processed with the QIAamp DNA Mini Kit (Cat. No. 51304) using the spin protocol of the manufacturer for DNA purification from blood or body fluids (Qiagen, 2020).

### Spectrophotometry

DNA concentration and purity were measured with a Multiskan SkyHigh spectrophotometer (Thermo Scientific) from the absorbance at 260 and 280 nm (Sambrook & Russell, 2001). Purity was assessed from the A_260_/A_280_ and A_260_/A_230_ ratios. A ratio A_260_/A_280_ of about 1.8 indicates minimal protein contamination, and A_260_/A_230_ of 1.8 or higher indicates low carry-over of chaotropic salts and polysaccharides.

### PCR amplification and gel electrophoresis

The mitochondrial hypervariable region 1 (HVR1) was amplified with the primers CAP-F (5′-CGTGTATGCAAGTACATTAC-3′) and CAP-R (5′-CTGATTAGTCATTAGTCCATC-3′) (Eurofins, India) described by Naderi et al. (2008). The template concentration was 30 ng µL^-1^. Each 20-µL reaction contained 2 U of Taq DNA polymerase (Takara Bio Inc., Cat. No. R001A), 1× Taq buffer with MgCl_2_, 0.125 mM dNTPs and 0.5 µM of each primer. Cycling in a ProFlex PCR System (Thermo Scientific) was 95 °C for 5 min, 35 cycles of 95 °C for 30 s, 60 °C for 30 s and 72 °C for 30 s, and 72 °C for 10 min. A no-template control (NTC) was included in every run. The products (5 µL) were resolved on a 1% agarose gel (UltraPure Agarose, Thermo Fisher, Cat. No. 16500500) stained with ethidium bromide (Sigma, Cat. No. E7637) in 1× TBE buffer (Tris–borate–EDTA) at 200 V, with a 100–1000 bp ladder (Thermo Fisher Scientific, Cat. No. 15628019) as size marker. Gels were imaged with the iBright Imaging System (Thermo Fisher Scientific), and band intensities were quantified by densitometry in ImageJ (v1.54, NIH).

### Statistical analysis

Because only three technical replicates from one straw were available, the inferential analysis is exploratory. DNA concentration (ng µL^-1^), A_260_/A_230_, A_260_/A_280_, normalised yield (ng µL^-1^ semen) and cost (INR per ng of DNA; INR = Indian rupees) were the dependent variables. Normality of residuals was tested with the Shapiro–Wilk test and homogeneity of variance with Levene’s test, separately for each variable. Normalised yield was the total DNA yield (ng) divided by the semen volume (µL), and cost per ng was the cost per sample divided by the total DNA yield, both calculated for each replicate. Residuals for concentration and normalised yield deviated from normality (Shapiro–Wilk W = 0.82, *p* = 0.016, and W = 0.74, *p* = 0.002, respectively), so the Kruskal–Wallis H test was the primary comparison for these variables, with Dunn’s test and Bonferroni correction for pairwise comparisons. Residuals for the purity ratios and for cost per ng were normal (Shapiro–Wilk *p* > 0.05), so one-way analysis of variance (ANOVA) with Tukey’s honest significant difference (HSD) test was used for them. The ANOVA for concentration and normalised yield is reported only as a secondary, descriptive comparison. The four protocols with three replicates (12 observations) entered the tests. Protocol B (n = 1) was excluded from all inferential models. Analyses were done in Python (v3.11) with SciPy v1.11.4 (Virtanen et al., 2020) and statsmodels v0.14.0 (Seabold & Perktold, 2010), and the data were handled with pandas v2.1.1 (McKinney, 2010). Results are given as mean ± sample standard deviation (SD).

## RESULTS AND DISCUSSION

### DNA yield

The five protocols differed markedly in DNA yield (Table 1; Figure 1). Modified Protocol 2 gave the highest concentrations in all three replicates (89.6, 47.8 and 30.4 ng µL^-1^; mean ± SD 55.9 ± 30.4 ng µL^-1^), followed by Kit B (15.5 ± 4.2 ng µL^-1^) and Modified Protocol 1 (14.8 ± 5.0 ng µL^-1^). The unmodified blood protocol (Protocol B) and the stool protocol (Protocol BS) gave the lowest concentrations (9.04 ng µL^-1^, n = 1, and 5.75 ± 0.35 ng µL^-1^, respectively). Expressed per µL of semen, the normalised yield of Modified Protocol 2 (55.9 ± 30.4 ng µL^-1^ semen) was about seven to ten times that of Modified Protocol 1 (5.9 ± 2.0), Kit B (7.8 ± 2.1) and Protocol BS (5.75 ± 0.35; Table 2). The Kruskal–Wallis test, the primary analysis, was significant for concentration (H = 9.36, *p* = 0.025), with Dunn’s test showing a difference only between Modified Protocol 2 and Protocol BS (*p* = 0.013). For normalised yield it was not significant (H = 6.90, *p* = 0.075), and no Dunn–Bonferroni comparison was significant (smallest *p* = 0.105, Modified Protocol 2 against Modified Protocol 1). The descriptive ANOVA of normalised yield gave F_3,8_ = 7.86 and *p* = 0.009, and Tukey’s HSD separated Modified Protocol 2 from each of the other three protocols (*p* = 0.016–0.020). Because the primary test was not significant, the samples were few and normality was violated, this parametric result is hypothesis-generating and not confirmatory. Variances were homogeneous (Levene’s test, *p* = 0.145). The large variation among replicates of Modified Protocol 2 (SD = 30.4 ng µL^-1^) probably reflects variation in the accessibility of sperm DNA among aliquots and emphasises the need for biological replication.

**Table 1.** Concentration, purity, total yield, processing time and cost of DNA extracted from frozen Black Bengal goat semen with five protocols.

| Protocol | Rep | Semen (μL) | DNA (ng μL<sup>-1</sup>) | A<sub>260</sub>/A<sub>230</sub> | A<sub>260</sub>/A<sub>280</sub> | Total yield (ng) | Time (min) | Cost (INR) |
| --- | --- | --- | --- | --- | --- | --- | --- | --- |
| Protocol B (Kit A, blood) | 1 | 250 | 9.04 | 0.601 | 1.27 | 904 | 40–45 | 300 |
| Modified Protocol 1 (Kit A) | 1 | 250 | 14.1 | 0.623 | 1.45 | 1410 | 40–45 | 336 |
| Modified Protocol 1 (Kit A) | 2 | 250 | 20.2 | 0.636 | 1.55 | 2020 | 40–45 | 336 |
| Modified Protocol 1 (Kit A) | 3 | 250 | 10.2 | 0.423 | 1.63 | 1020 | 40–45 | 336 |
| Modified Protocol 2 (Kit A) | 1 | 100 | 89.6 | 2.25 | 1.79 | 8960 | 190–200 | 336 |
| Modified Protocol 2 (Kit A) | 2 | 100 | 47.8 | 2.07 | 1.77 | 4780 | 190–200 | 336 |
| Modified Protocol 2 (Kit A) | 3 | 100 | 30.4 | 1.85 | 1.77 | 3040 | 190–200 | 336 |
| Kit B (silica membrane) | 1 | 200 | 17.8 | 1.52 | 1.84 | 1780 | 50–60 | 470 |
| Kit B (silica membrane) | 2 | 200 | 10.6 | 1.57 | 1.83 | 1060 | 50–60 | 470 |
| Kit B (silica membrane) | 3 | 200 | 18.1 | 0.728 | 1.32 | 1810 | 50–60 | 470 |
| Protocol BS (Kit A-S, stool) | 1 | 200 | 5.78 | 0.104 | 1.23 | 1156 | 45–50 | 300 |
| Protocol BS (Kit A-S, stool) | 2 | 200 | 5.39 | 0.074 | 1.25 | 1078 | 45–50 | 300 |
| Protocol BS (Kit A-S, stool) | 3 | 200 | 6.08 | 0.145 | 1.03 | 1216 | 45–50 | 300 |

Kit A = QIAamp DNA Mini Kit (Qiagen); Kit A-S = QIAamp DNA Mini Kit, stool protocol; Kit B = PureLink Genomic DNA Mini Kit (Invitrogen). Rep. = technical replicate from one frozen semen straw (one extraction for Protocol B). DNA was eluted in 100 μL (Kit A, Kit B) or 200 μL (Kit A-S). INR = Indian rupees (USD 3.16 for INR 300, 3.54 for INR 336 and 4.95 for INR 470).

**Table 2.** Normalised yield, purity and cost of the five DNA extraction protocols.

| Protocol | Normalised yield (ng $\mu\text{L}^{-1}$ semen) | $A_{260}/A_{230}$ | $A_{260}/A_{280}$ | Cost per sample (INR) | Cost per ng DNA (INR) |
| --- | --- | --- | --- | --- | --- |
| Protocol B (Kit A, blood) | 3.62 (n = 1) | 0.60 (n = 1) | 1.27 (n = 1) | 300 | 0.332 (n = 1) |
| Modified Protocol 1 (Kit A) | $5.93 \pm 2.02$ | $0.56 \pm 0.12$ | $1.54 \pm 0.09$ | 336 | $0.245 \pm 0.082$ |
| Modified Protocol 2 (Kit A) | $55.93 \pm 30.43$ | $2.06 \pm 0.20$ | $1.78 \pm 0.01$ | 336 | $0.073 \pm 0.037$ |
| Kit B (silica membrane) | $7.75 \pm 2.12$ | $1.27 \pm 0.47$ | $1.66 \pm 0.30$ | 470 | $0.322 \pm 0.105$ |
| Protocol BS (Kit A-S, stool) | $5.75 \pm 0.35$ | $0.11 \pm 0.04$ | $1.17 \pm 0.12$ | 300 | $0.262 \pm 0.016$ |

**Figure 1.**
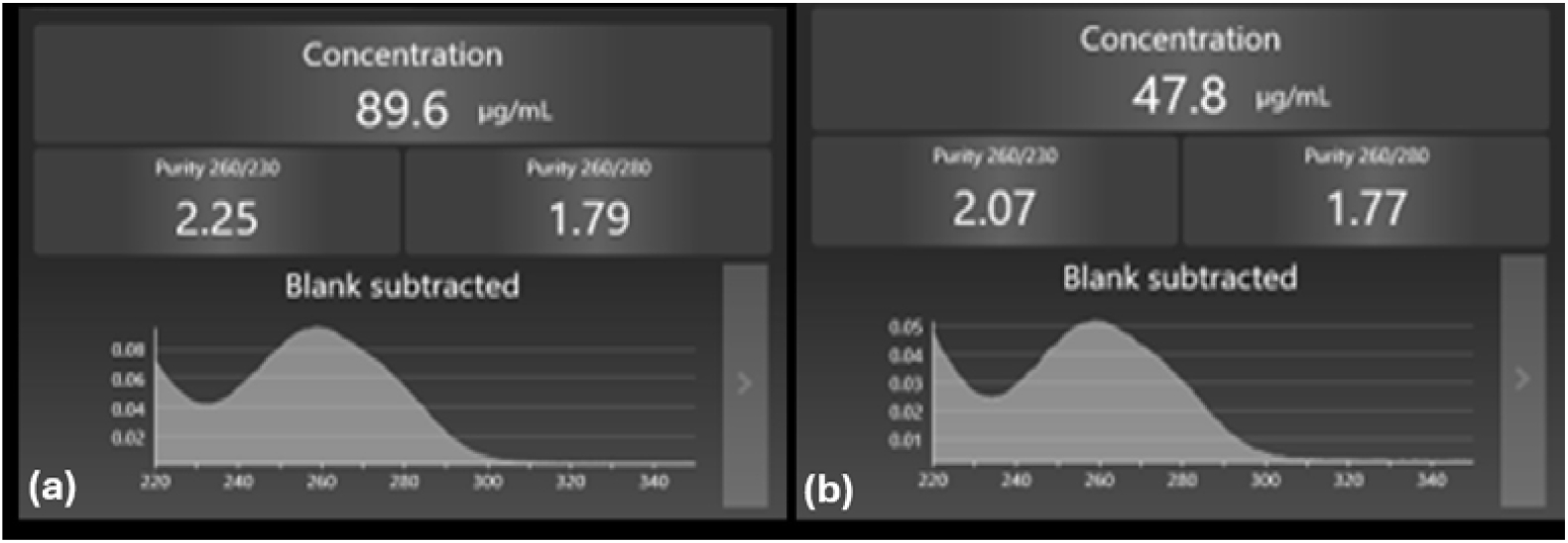
Spectrophotometric readouts (Multiskan SkyHigh) of DNA extracted with Modified Protocol 2 (Kit A). (a) Replicate 1: 89.6 ng µL^-1^, A_260_/A_280_ = 1.79, A_260_/A_230_ = 2.25. (b) Replicate 2: 47.8 ng µL^-1^, A_260_/A_280_ = 1.77, A_260_/A_230_ = 2.07. Blank-subtracted absorbance spectra, 220–350 nm.

### Mechanism of the stepwise digestion

Modified Protocol 2 was the most effective method. Its advantage is probably due to the stepwise proteinase K strategy. Disulphide bonds are central to the compaction of bull sperm chromatin (Hutchison et al., 2017), and the nucleoprotamine structure can be partly unpacked chemically with sodium dodecyl sulphate and dithiothreitol (DTT) (Filho et al., 2015). The first 10-µL aliquot of enzyme presumably digests protamine-associated proteins in part, and the second acts on the newly exposed chromatin, which overcomes the enzyme saturation that limits protocols with a single addition. The total digestion time (3 h) was longer than that of Modified Protocol 1 (2 h) and of the unmodified kit protocol (about 30 min). The smaller starting volume (100 µL against 200–250 µL) may also have reduced the inhibitor load on the spin column and improved binding to the silica membrane. A reducing agent such as 1–5 mM DTT or 0.1% β-mercaptoethanol before digestion could break the disulphide cross-links directly and might raise the yield further, as reported in related forensic and livestock work (Vuichard et al., 2011; Silva et al., 2014). Exposure to 2 mM DTT also improved access to the chromatin of human spermatozoa in a DNA damage assay (Mitchell et al., 2011). This should be tested in future work.

### DNA purity

Modified Protocol 2 and Kit B both gave A_260_/A_280_ values close to the optimum of about 1.8 (1.78 and 1.66, respectively; Table 2). Modified Protocol 2 also gave the highest mean A_260_/A_230_ (2.06 ± 0.20), which indicates minimal carry-over of chaotropic salts or polysaccharides. This ratio is sensitive to residual salts and seminal plasma components and is often a better predictor of PCR inhibition than A_260_/A_280_ alone (Wilson, 1997). The stool protocol (Protocol BS) gave much lower A_260_/A_280_ values (1.17 ± 0.12), which indicates substantial protein contamination that can inhibit Taq polymerase and other enzymes (Wilson, 1997). One-way ANOVA showed significant differences among the protocols for A_260_/A_280_ (F_3,8_ = 7.47, *p* = 0.011) and for A_260_/A_230_ (F_3,8_ = 31.3, *p* < 0.001). Tukey’s HSD showed that A_260_/A_280_ of Protocol BS was lower than that of Modified Protocol 2 (*p* = 0.009) and Kit B (*p* = 0.028), whereas Modified Protocol 2 and Kit B did not differ. A_260_/A_230_ of Modified Protocol 2 was higher than that of every other replicated protocol (*p* ≤ 0.027). Kit B gave acceptable purity but a lower yield, in line with the known trade-off between purity and yield in pre-packed silica column kits (Kumari et al., 2019). It is adequate for PCR but may be limiting where more template DNA is needed, as in whole-genome sequencing.

### DNA integrity and PCR compatibility

Gel electrophoresis showed distinct high-molecular-weight bands for all protocols (Figure 2). Band intensity was greatest for Modified Protocol 2 (lanes 3–6), as confirmed by densitometry. DNA from Kit B (lanes 7–8) amplified successfully, but with weaker bands, in line with its lower yield (Figure 3). The no-template controls gave no amplification. The stool protocol (lanes 9–13) and the unmodified blood protocol (lane 1) gave weak or no amplification, in line with their low yield and poor purity.

**Figure 2.**
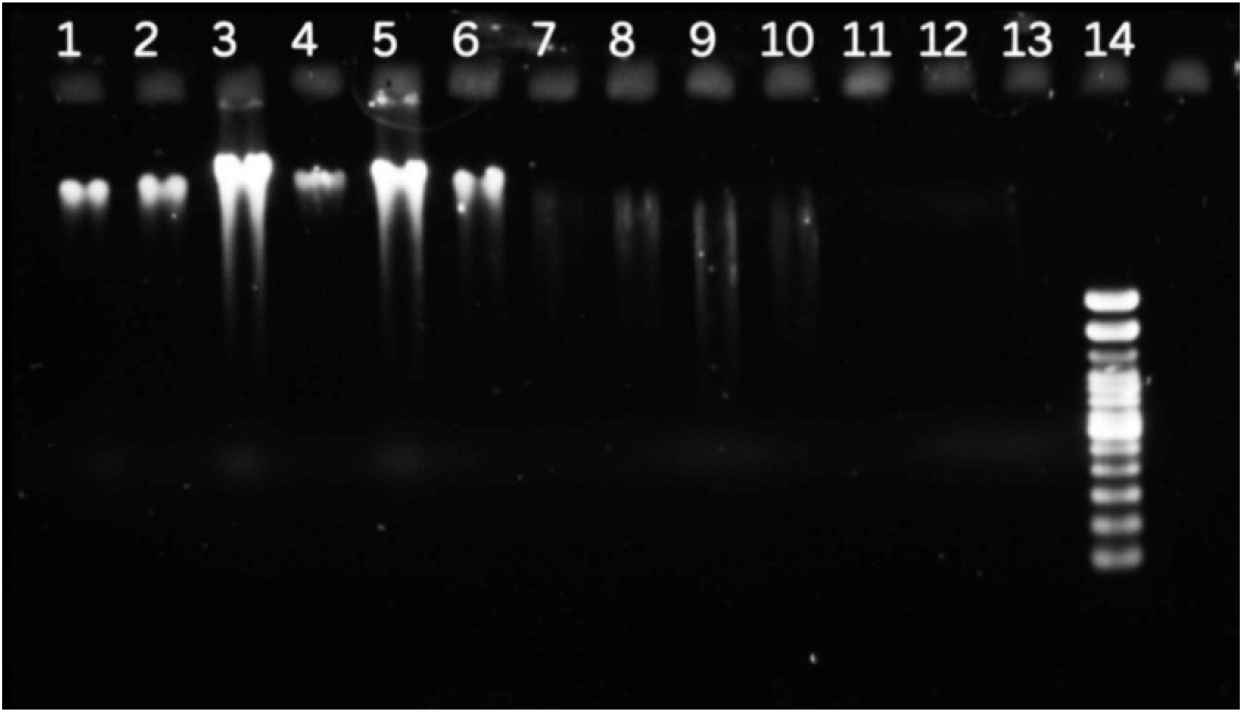
Agarose gel (1%) of genomic DNA extracted with five protocols. Lane 1: Protocol B (Kit A, blood). Lane 2: Modified Protocol 1 (Kit A). Lanes 3–6: Modified Protocol 2 (Kit A). Lanes 7–8: Kit B (silica membrane). Lanes 9–13: Protocol BS (Kit A-S, stool). Lane 14: 100–1000 bp ladder.

**Figure 3.**
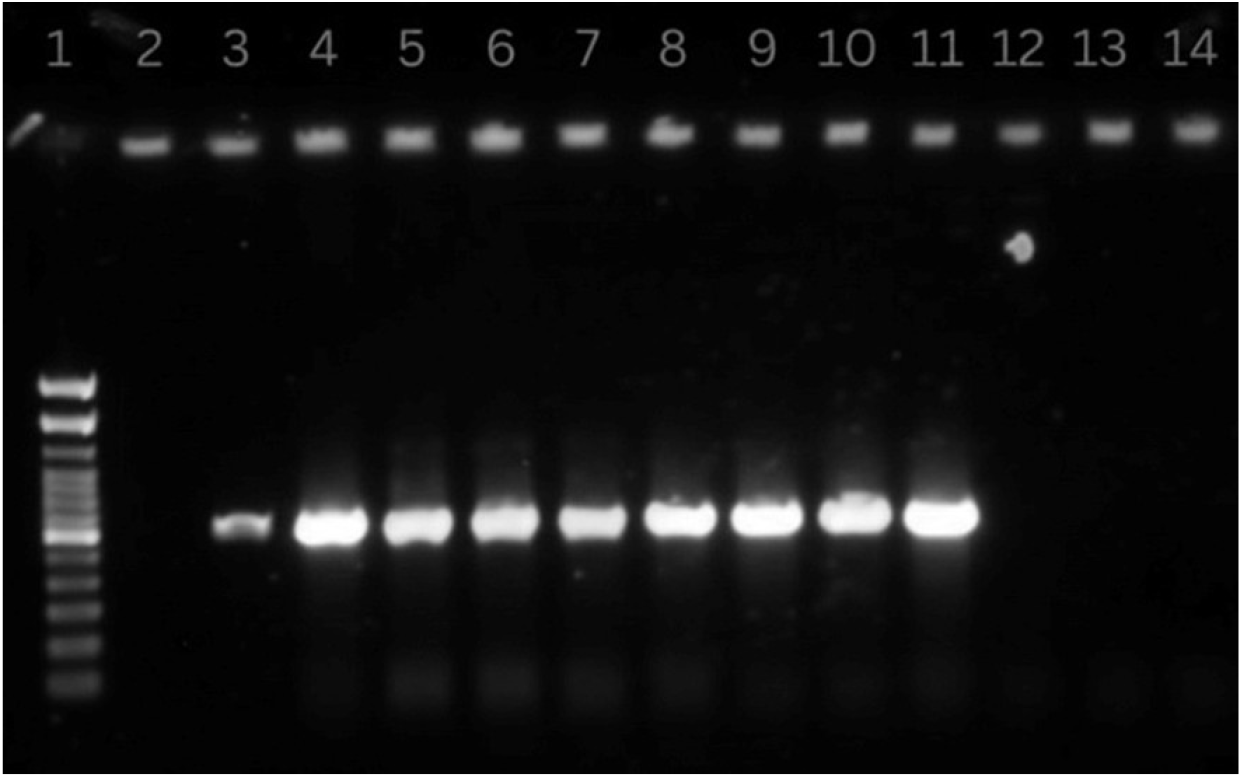
Agarose gel (1%) of PCR products of the mitochondrial hypervariable region 1 (HVR1, about 430 bp). Lane 1: 100–1000 bp ladder. Lane 2: Modified Protocol 1 (Kit A). Lanes 3–6: Modified Protocol 2 (Kit A). Lanes 7–8: Kit B (silica membrane). Lanes 9–14: Protocol BS (Kit A-S, stool). The no-template control showed no amplification (not shown).

### Cost efficiency and limitations

Modified Protocol 2 cost INR 336 per sample, which was lower than Kit B (INR 470). Because of its higher yield, the cost per ng of DNA was INR 0.073 ± 0.037 for Modified Protocol 2, against 0.322 ± 0.105 for Kit B, 0.245 ± 0.082 for Modified Protocol 1 and 0.262 ± 0.016 for Protocol BS (Table 2); Protocol B gave 0.332 (n = 1). Modified Protocol 2 was about fourfold cheaper per ng than Kit B (Tukey’s HSD, *p* = 0.010) and cheaper than Protocol BS (*p* = 0.042), but not significantly cheaper than Modified Protocol 1 (*p* = 0.063; ANOVA F_3,8_ = 7.14, *p* = 0.012). Its hands-on time was longer (190–200 min against 50–60 min for Kit B), but the higher yield and better purity give a better balance between resources and the usefulness of the DNA for routine genomic studies. The main limitation is the use of technical replicates from a single straw, which prevents any inference about variation among animals. The large variation among replicates of Modified Protocol 2 (SD = 30.4 ng µL^-1^) cannot be separated from biological variation without biological replication. Modified Protocol 2 also differed from the other protocols in both the proteinase K scheme (stepwise against a single addition) and the starting volume (100 µL against 200–250 µL), so their separate contributions cannot be determined and a factorial design is needed. DNA integrity was assessed visually and by densitometry, and the DNA Integrity Number (DIN) from a fragment analyser would give more rigorous data for sequencing applications. Only Black Bengal goats were included, and amplicon identity was inferred from fragment size and not confirmed by Sanger sequencing.

## CONCLUSION

Five DNA extraction protocols for goat semen were compared using three technical replicates from one Black Bengal buck semen straw. Modified Protocol 2, a re-optimised spin-column protocol with stepwise proteinase K digestion over 3 h, gave the highest normalised yield (55.9 ± 30.4 ng µL^-1^ semen), the best purity ratios (A_260_/A_230_ about 2.06 and A_260_/A_280_ about 1.78) and the lowest cost per ng of DNA (INR 0.073 ± 0.037). Its extracts, like those of Kit B, amplified in the HVR1 PCR, whereas the blood and stool protocols of Kit A gave weak or no amplification. The findings are preliminary. Confirmation with biological replicates from at least five bucks, including other breeds, is needed before Modified Protocol 2 can be recommended for routine large-scale genomic studies, and reducing agent supplementation and DIN measurement should be tested.

## ACKNOWLEDGEMENT

The authors thank the Chief Executive Officer of Paschimbanga Go Sampad Bikash Sanstha (PBGSBS), the Managing Director of the West Bengal Livestock Development Corporation Limited (WBLDCL) and the Directorate of Animal Husbandry and Veterinary Services, Government of West Bengal, for guidance and institutional support. Gemini was used to modify the citation style. Quill Bot was used to reformat the manuscript according to the journal template. The authors checked the content and take full responsibility for it.

## CONFLICT OF INTEREST

The authors declare that they have no conflicts of interest.

## AUTHOR/S DECLARATION

- We confirm that all Figures and Tables in the manuscript are original to us. Additionally, any Figures and images that do not belong to us have been incorporated with the required permissions for re-publication, which are included with the manuscript.
- Ethical clearance and animal welfare: frozen semen straws from routine artificial insemination archive stock were used; no live animals were handled and no experimental procedures were performed on animals, so Institutional Animal Ethics Committee approval was not applicable.
- Funds: This research did not receive any specific grant from funding agencies in the public, commercial or not-for-profit sectors.

## AUTHOR’S CONTRIBUTION STATEMENT

Sharadindu Shil: conceptualization, methodology, original draft. Ankur Singha: investigation, data curation. Soumita Paul, Ashish Khatua, Shankari Prasad Dutta and Tarpan Mondal: investigation, laboratory experiments. Jayanta Chowdhury and Ayan Mukherjee: resource management. Anusuya Debnath and Ajoy Mandal: formal analysis, supervision, review and editing. Author Contribution Statement form to be completed.

## Notes

### Competing Interest Statement

The authors have declared no competing interest.

